# Pathways of DNA unlinking: A story of stepwise simplification

**DOI:** 10.1101/188722

**Authors:** Robert Stolz, Masaaki Yoshida, Reuben Brasher, Michelle Flanner, Kai Ishihara, David J. Sherratt, Koya Shimokawa, Mariel Vazquez

**Affiliations:** Department of Microbiology and Molecular Genetics, University of California Davis, Davis, USA; Department of Mathematics, Saitama University, Saitama, Japan; Microsoft, San Francisco, USA; Faculty of Education, Yamaguchi University, Yamaguchi, Japan; Department of Biochemistry, University of Oxford, Oxford, UK; Department of Mathematics, University of California Davis, Davis, USA; Takasaki City Office, 35-1 Takamatsu-cho, Takasaki, Japan

## Abstract

In ***Escherichia coli*** DNA replication yields interlinked chromosomes. Controlling topological changes associated with replication and returning the newly replicated chromosomes to an unlinked monomeric state is essential to cell survival. In the absence of the topoisomerase topoIV, the site-specific recombination complex XerCD-*dif*-FtsK can remove replication links by local reconnection. We previously showed mathematically that there is a unique minimal pathway of unlinking replication links by reconnection while stepwise reducing the topological complexity. However, the possibility that reconnection preserves or increases topological complexity is biologically plausible. In this case, are there other unlinking pathways? Which is the most probable? We consider these questions in an analytical and numerical study of minimal unlinking pathways. We use a Markov Chain Monte Carlo algorithm with Multiple Markov Chain sampling to model local reconnection on 491 different substrate topologies, 166 knots and 325 links, and distinguish between pathways connecting a total of 881 different topologies. We conclude that the minimal pathway of unlinking replication links that was found under more stringent assumptions is the most probable. We also present exact results on unlinking a 6-crossing replication link. These results point to a general process of topology simplification by local reconnection, with applications going beyond DNA.

## 1 Introduction

Flexible circular chains appear often in nature, from microscopic DNA plasmids to macroscopic loops in solar corona. Such chains entrap rich geometrical and topological complexity which can give insight into the processes underlying their formation or modification. Knotted and interlinked states often coincide with higher energy states in physical systems and are usually undesired. Topology-simplifying reconnection processes involving one or two cleavages are observed. Examples in biology include the action of type II topoisomerases and of site-specific recombinases. Type II topoisomerases bind to two segments of double-stranded DNA, cleave one of the segments, transport the other through the break (*strand-passage*) and reseal the break. Site-specific recombinases bind to two specific sites (short segments of double-stranded DNA), introduce a double-stranded break on each site, recombine the ends and reseal the breaks. The action of recombination enzymes is a local reconnection event. We here investigate pathways of unlinking of newly replicated DNA links by local reconnection. The results presented, and the numerical methods proposed are not restricted to the biological example and are applicable to any local reconnection process.

In genetics, the observation of topological links dates back to studies in plants in the 1930s. In a study of chromosomal variation in *Crepis tectorum*, M. Navashin observed ring chromosomes, noting “in one case, the two daughter strands composing a normal chromosome failed to separate". Navashin reported on a metaphase involving four rings, two of which were “united in the fashion of chain links," thus documenting the appearance of two newly replicated circular chromosomes forming a singly-linked catenane, or 2-crossing link.^1^ In her study of ring chromosomes in maize, Barbara McClintock observed the accumulation of several rings in the same cell and hypothesized that “lack of uniformity in the splitting plane could give rise to a double sized ring with two insertion regions or cause split halves of the ring to become interlocked", thus introducing the ideas of chromosome dimers and links (also called catenanes).^2^ Three decades later, DNA links were studied *in vitro* via random cyclization of circular DNA in the presence of an excess of DNA circles^3^ and, in 1980 interlinked dimers formed by nicked newly replicated 5.2kb circular dsDNA mini chromosomes from SV40 were observed by electron microscopy.^4^ The mechanisms of replication and segregation of circular DNA predict products that can be topologically characterized as right-hand (RH) 2m-crossing torus links with parallel sites, which we here refer to as parallel 2m-cats (denoted mathematically as parallel 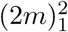 or *T*(2,2*m*)_*p*_).^5^ These topological forms were confirmed by characterizing the linked replication intermediates that accumulate in topoIV mutants^6^ (Fig. 1(A)). Sogo *et al.*^8^ hypothesized that catenanes appeared as replication intermediates of bacteriophage λ DNA and observed that, in order to secure proper segregation of circular chromosomes at cell division, the linking number of the two newly replicated molecules must be reduced to zero. However, the topology of a circular double-stranded (ds)DNA molecule is insensitive to any manipulation that does not allow a double-stranded break.^5^ Nicking of a single DNA strand, however extensive, is insufficient to unlink two newly replicated DNA circles unless pre-existing nicks are present along the second strand. The type II topoisomerase topoIV is a major decatenase in *E. coli.*^6,9^ Grainge *et al.* showed that in the absence of topoIV, the XerCD-dif-FtsK molecular machine can act *in vivo* to separate two interlinked, newly replicated chromosomes.^10^ The XerCD complex consists of the site-specific tyrosine recombinases XerC and XerD. The *dif* site is a 28bp long recombination site located within the terminus region of the E. coli chromosome. FtsK is a powerful translocase that assembles at the division septum, where it activates XerCD-dif recombination. Their experimental data suggested a gradual reduction in topological complexity of the substrates, which were RH 2m-cats with parallel *dif* sites.^10^ The proposed unlinking pathway, through which the enzymes unlink the replication links in a step-wise fashion is illustrated in Fig. 1A. In the figure, each closed curve represents a circular dsDNA molecule. The components of a two-component link represent two newly replicated DNA chains.

**Figure 1:**
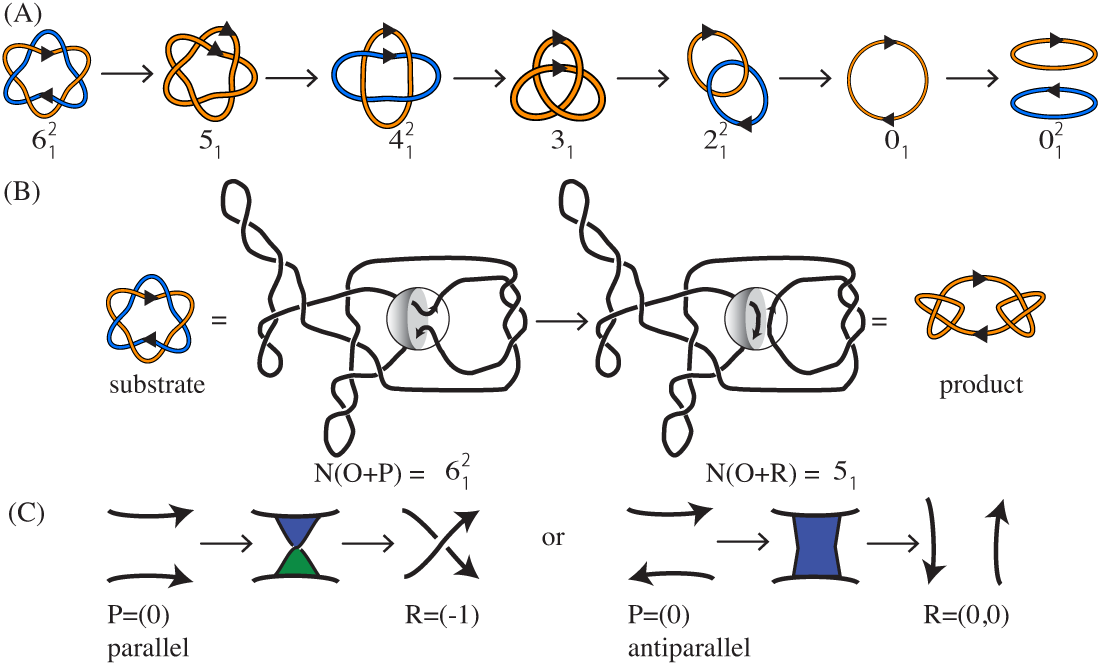
(A) Under the assumption that each reconnection step strictly reduces the number of crossings of the substrate, in Shimokawa *et al.*^7^ we showed that there is a unique unlinking pathway starting at a 2*m*-crossing replication link. In *E. coli* a replication link is a 2*m*-cat with parallel *dif* sites,^6^ and this pathway predicts the first product to be a (2*m* – 1) i knot with two *dif* sites in direct repeats. Two sites along a knotted chain are in *direct repeats* if they induce the same orientation into the knot. Replication links are 2*m*-crossing right-handed torus links with parallel sites (mathematical notation: 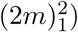. The pathway in the figure illustrates, for *m* = 6, the only unlinking pathway starting at the parallel 2*m*-cat under the assumption that each reconnection step strictly reduces the minimal crossing number. All the intermediate topologies are torus links 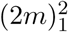 or torus knots (2*m* – 1)_1_ with two reconnection sites in direct repeats as in the figure. (B) One reconnection step: here the cleavage regions of the reconnection sites on a 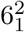 link are brought together to form a synapse (shown as a ball enclosing two strings). The synapse is modeled mathematically as a 2-string tangle. In the case of XerCD site-specific recombination, the strings in the tangle contain the core regions of the *dif* sites (indicated by two arrows in a tangle *P* representing two very short segments of double-stranded DNA which physically behave as two almost straight strings) and any bound DNA which does not change during recombination (gray shaded regio*n*). Any interesting geometrical or topological complexity of the substrate is captured mathematically as an outside tangle *O* that remains constant during reconnection. Before strand cleavage, the substrate is modeled by the tangle equation 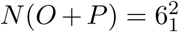. The local reconnection is modeled by tangle surgery where *P* is replaced with *R*, yielding a product represented as *N*(*O* + *R*) = *K*, where *K* is a knot with two directly repeated sites. (C) Local reconnection is a simple event which can be modeled as a band surgery, where *P* = (0) is replaced with a tangle *R* = (*w*, 0) enclosing a vertical row of *w* twists, for some integer *w*. The rational tangle notation (or Conway notatio*n*) for such vertical tangle is *R* = (*w*, 0). In the case when *w* = ±1 the notation simplifies to *R* = (±1). In the simplest cases, *P* = (0) with sites in parallel alignment goes to *R* = (±1), and *P* = (0) with sites in anti-parallel alignment goes to *R* = (0,0) as illustrated in the figure.

A rigorous mathematical analysis of the recombination experiments of Grainge *et al*.^10^ showed that at least 2*m* steps are needed in order to unlink any RH 2*m*-cat with parallel sites.^7^ This result relied simply on the assumption that the XerCD tetramer binds the two *dif* sites and that a simple cut-reconnect-paste reaction ensues (Fig. 1C). If the shortest pathway of unlinking a 2*m*-crossing replication link has exactly 2*m* steps, it is natural to ask how many such pathways exist and whether some are more likely than others. Under the assumption that each step strictly reduces the topological complexity of its substrate (as measured by minimal crossing number), Shimokawa et al.^7^ showed that the only possible pathway of unlinking a 2*m*-crossing replication link is that in Fig. 1A. Using tangle calculus, they proposed a 3-dimensional topological mechanism to take the parallel 2*m*-cat to the unlink. This mechanism incorporates three solutions obtained by tangle calculus at each step of the process, and the last three steps are fully characterized. The results in Shimokawa *et al.*^7^ provide unprecedented detail in the study of the topological mechanism of DNA unlinking by site-specific recombination. Going beyond the original problem of unlinking newly replicated circular chromosomes, these results apply to any reconnection event that can be modeled using tangles as in Fig. 1. For example, the same unlinking pathway proposed for DNA links under site-specific recombination has been observed during reconnection events in physical fields such as vortices in fluid flow.^11–13^ Further mathematical research on this subject can be found in the literature.^14–18^

Successful unlinking by XerCD-FtsK of newly replicated plasmids containing *dif* sites was shown in.^10^ Quantification of these data gave weak justification to the assumption of stepwise reduction in complexity during the unlinking reaction.^7^ As can be seen in Fig. 2, the gel quantification clearly illustrates the reduction of replication links by XerCD-FtsK site-specific recombination at *dif* sites. However, because of the complexity of the data, in order to confirm stepwise reduction one would need to repeat the time course experiments^10^ for each individual topology. This motivates the current work where we remove the assumption of stepwise decrease in complexity, and design mathematical and numerical methods to assess the different unlinking pathways and the identification of the most probable ones. We ask whether there are other minimal unlinking pathways and hypothesize that the minimal pathway previously proposed^7,10,19^ and illustrated in Fig. 1A is the most likely among all the possible minimal pathways that arise. First, we allow the complexity of the products to decrease or remain the same at each step of the reaction. We provide analytical proof that there are exactly nine minimal pathways of unlinking a parallel 6-cat; many of the resulting transitions are fully characterized. Characterizing minimal pathways of unlinking by local reconnection and resolving the topological mechanisms involved are problems of high theoretical complexity since the number of possibilities quickly increases with the number of crossings of the substrate. Likewise, characterizing the topological mechanism(s) taking a link *L*_*i*_ to a knot *K*_*j*_ is equivalent to characterizing all band surgeries between *L*_*i*_ and *K*_*j*_ (see Fig. 1C).

**Figure 2:**
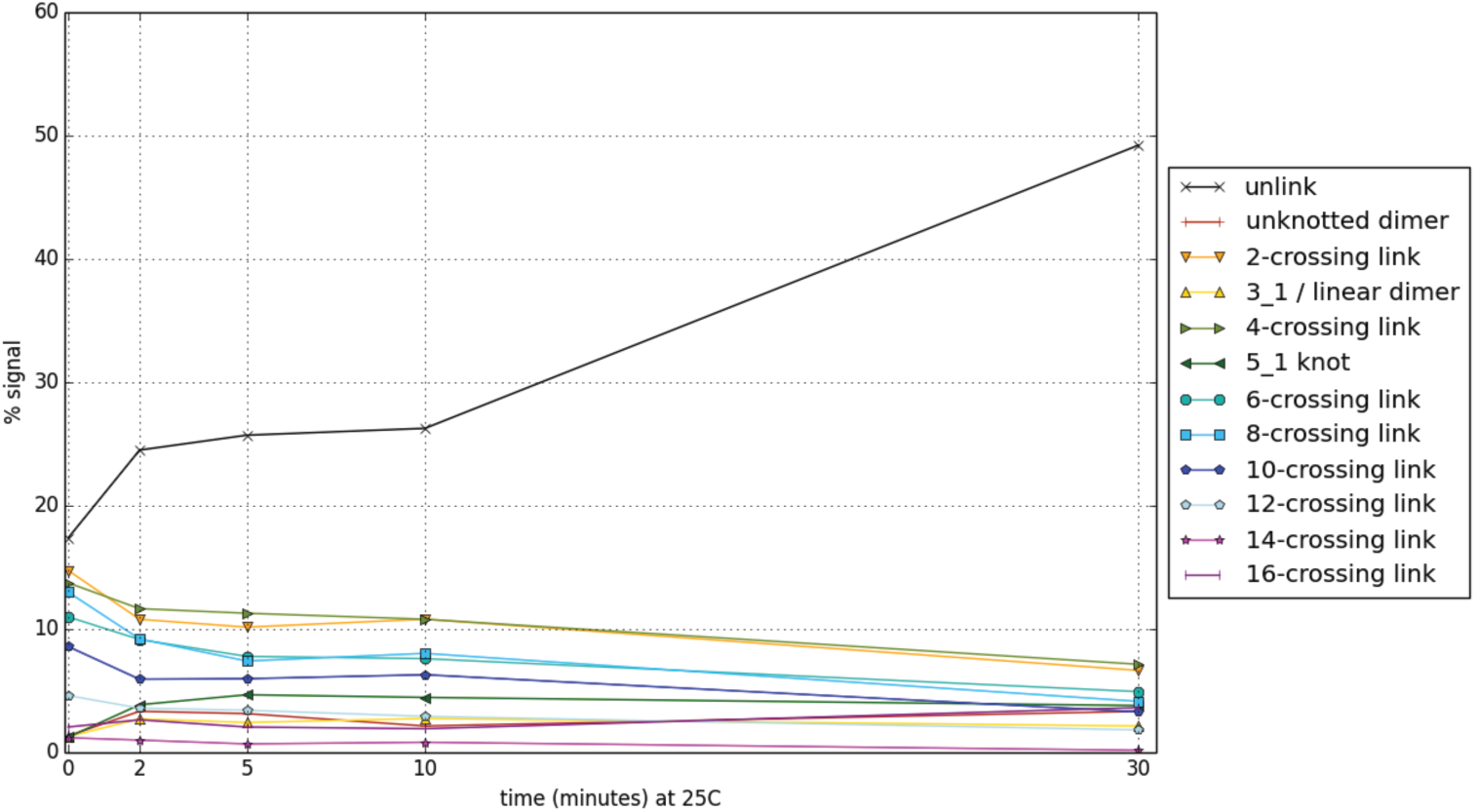
Quantification of the time-course experiments.^10^ The gel presented in Fig. 1B in Grainge *et al.*^10^ showed a time course of unlinking by XerCD-*dif*-FtsK50C at 25^o^C of newly replicated plasmids containing *dif* sites. Line scans of the gel were previously published.^7^ In this figure each topological class is shown as a separate series of points with linear interpolation. The caption assumes the bands observed correspond to the topologies expected from a substrate composed of replication links, i.e. 2*m*-crossing links (e.g. 2*m*-cats), and some of the corresponding knotted intermediates (open circle or 0_1_,3_1_, 5_1_). "Unlink" corresponds to the two unlinked components in monomeric state (topology type 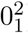), and "Unknot" corresponds to the dimeric unknot (0_1_). The quantification clearly illustrates the reduction of replication links by XerCD-FtsK site-specific recombination at *dif* sites. The complexity of the data is also evident, with the relative proportions of all the different topologies fluctuating from one step to the next, thus obscuring the signal.

In order to discriminate between different minimal unlinking pathways for a given substrate and to extend the study to higher crossing numbers, we eliminate the complexity assumption and develop a Monte Carlo method to simulate local reconnection events. The method can be applied to a substrate with any topology, allows products of varying topological complexity, and facilitates the rigorous quantification of the transition probabilities along each obtained pathway. Using this method we embark on a numerical study relevant to unlinking of DNA replication links by site-specific recombination a *dif* sites. More specifically, we restrict the numerical study to knotted chains of fixed length with two reconnection sites (representing the *dif* sites) that are evenly spaced along the chain, and linked chains consisting of the union of two circles of same length with one reconnection site in each component. Details on the numerical experiments can be found in the Numerical Methods section and in the Supplementary Methods online.

The computational approach provides a rigorous means to discriminate between mathematically equivalent unlinking pathways. The combination of the mathematical and computational studies provides strong quantitative support for the hypothesis that the unlinking pathway from Fig. 1A is the most likely, even under the weakened assumptions.

**Nomenclature for knots and links** It is important at the outset to say a word about the naming convention used for the knots and links which arise in this study (490 knots and 391 two-component links). A local reconnection event on a two component link with one cleavage site in each component yields a knotted chain with two sites in direct repeats (*cf.* Fig. 1A). Rolfsen’s Knot Table^20^ summarizes the knot nomenclature used in the mathematics community, which was not intended to distinguish between mirror images nor between oriented links, an important consideration when dealing with circular DNA and other biopolymers. Chirality is relevant, and indeed crucial, to characterize biological and chemical compounds. In this paper, we use the writhe-based knot nomenclature proposed in Brasher et al.^21^ The *writhe* is a geometrical invariant that provides a measure of a chain’s entanglement complexity and chirality. It is computed analytically using a Gauss double integral and can be estimated numerically by taking the average of the writhe of a planar diagram taken over all projection directions (*the projected writhe*). The *mean writhe* of a knot *K* refers to the average of the writhes of all knotted chains of type *K*. Numerically this is estimated by averaging over a sufficiently large, randomly generated ensemble of conformations of type *K*. A representative of a chiral pair is chosen based on its mean writhe.^21^ We extend this nomenclature to the 2-component links depicted in Fig. 3. For prime 2-component links with 9 or more crossings we use the default notation from Knotplot.^22^ For more details and a comparison with other published nomenclature for links refer to the Supplementary Methods and to Supplementary Fig. S5 online.

**Figure 3:**
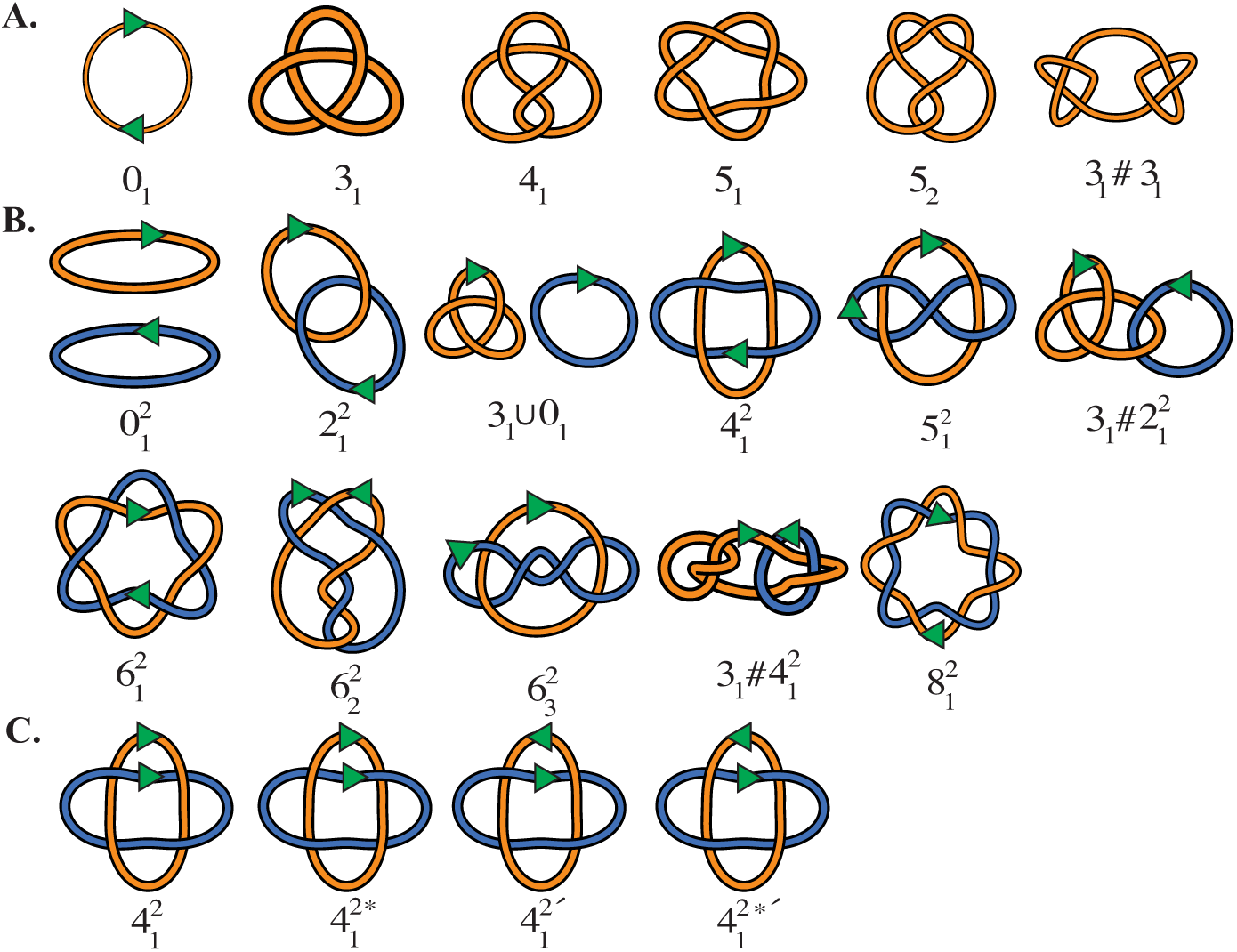
(A) Illustration of some of the knots relevant to the present study and their nomenclature. The chirality is consistent with that in Brasher *et al.*^21^ The green arrows along the unknot 0_1_ represent the two reconnection sites. The sites shown are equidistant and in direct repeats. A complete table of prime knots with up to 10 crossings and information on how they compare to those in Rolfsen^20^ can be obtained from the authors upon request. (B) Nomenclature for two component links relevant to the present study. The green arrows represent the reconnection sites, which confer an orientation to each link component. The nomenclature is described in the Supplementary Methods and in Supplementary Fig. S5 online. For 2-component links with 9 or more crossings we revert to the default Knotplot naming convention. (C) The four possible combinations of chirality and orientation for the 4-crossing torus link. A comparison between the nomenclature used in this paper and that in Rolfsen^20^ and in works by Kanenobu^23,24^ is included in Supplementary Fig. S5 online. Arrows indicate the relative orientations of the sites.

## 2 Results

### 2.1 There are exactly 9 shortest pathways to unlink the 6-cat that do not increase substrate complexity

We consider an event where two oriented sites come together and undergo cleavage followed by reconnection. If the substrate is a single circle, then the oriented sites are in direct repeat, *i.e.* they induce the same orientation into the circle. If the substrate consists of two circular chains, then there is one site in each chain. Note that such an event always changes the topology of the substrate: reconnection between two sites in separate components of a link yields a knot with two sites in direct repeats, and reconnection on a knot with two directly repeated sites yields a 2-component link with one site in each component. The reconnection event is modeled as a system of tangle equations as described in Fig. 1(B). In the context of DNA unlinking, as in Shimokawa *et al.*,^7^ we model dsDNA as a curve defined by the axis of the DNA double helix, and the synapse formed by the enzymes bound to the core regions of the *dif* recombination sites as the 2-string tangle P. Reconnection changes *P* into *R*. If we assume that each reconnection is modeled as a coherent band surgery, *i.e. P* = (0) and *R* = (*w*, 0) for some integer *w*, then any minimal pathway to unlink an n-crossing torus link with parallel sites (e.g. 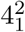 or 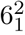) has exactly *n* steps. Furthermore, if each reconnection step is assumed to strictly reduce the complexity of its substrate, then the minimal pathway is unique: *i.e.* RH 2*m*-cat, RH (2,2*m* – 1)-torus knot, RH (2*m* – 2)-cat, · · ·, RH trefoil, Hopf link, trivial knot, trivial link. Fig. 1A illustrates the 6-cat case. Since the experimental data^10^ only gives weak support to the assumption that the complexity goes strictly down at each step of the reaction (Fig. 2), we here examine the case where no reconnection step increases the number of crossings and provide analytical characterization of all shortest pathways from the 6-cat to the unlink.

#### Assumption 1.

Consider a reconnection pathway from a parallel RH 2mcat to the unlink. Assume that each product along the pathway is a knot or a 2-component link, that the pathway is shortest, and that no reconnection event increases the number of crossings of its substrate.

Recall that any shortest reconnection pathway from (2*m*)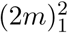 to the unlink has exactly 2*m* steps.^7^ In Theorem 2 we show that there are exactly nine unlinking pathways satisfying Assumption 1.

#### Theorem 2.

A pathway from the parallel RH 6-cat that satisfies Assumption 1 is one of the 9 shown in Fig. 4.

**Figure 4:**
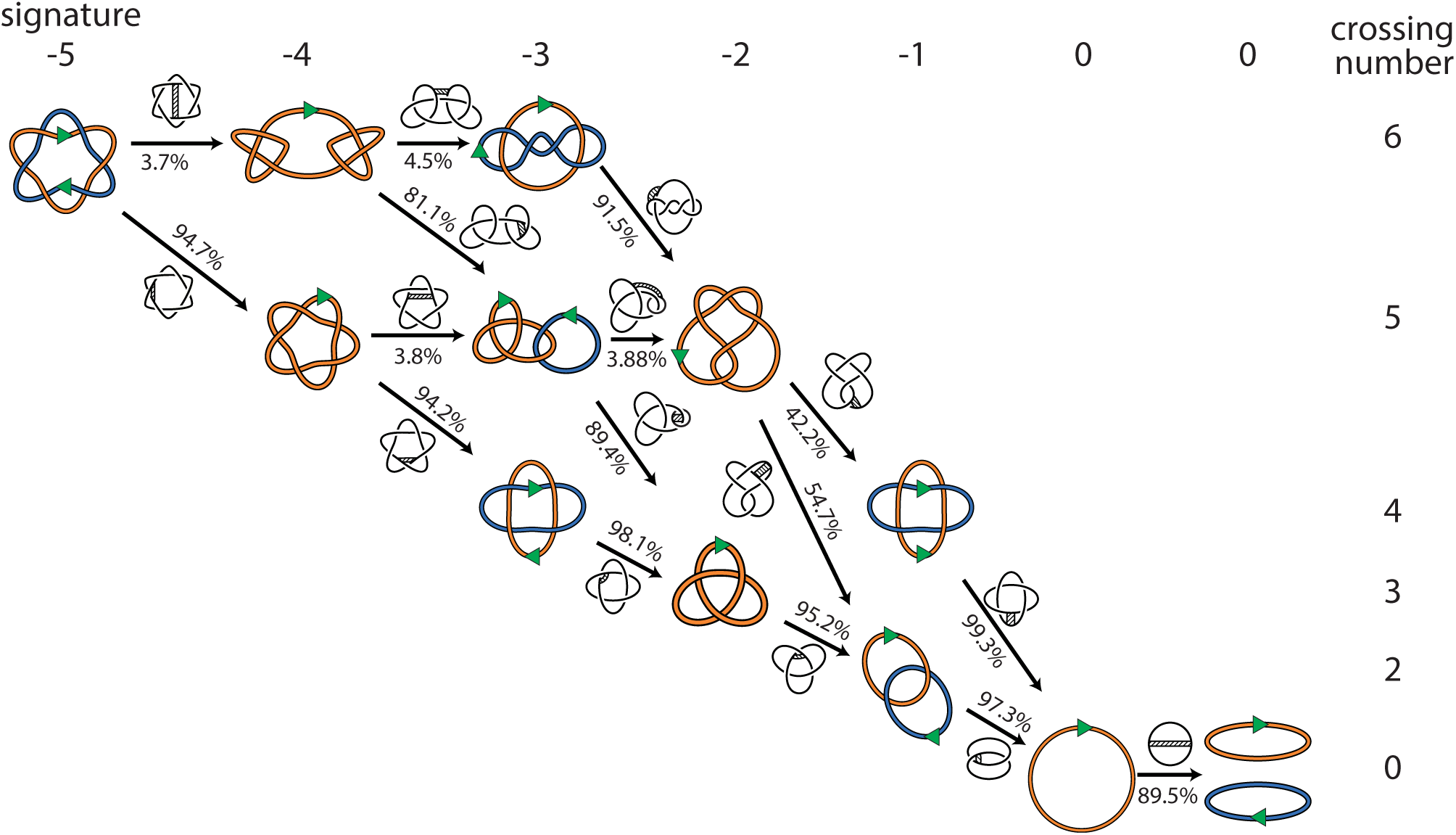
The substrate at the top left corner is the link 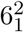 with two reconnection sites in parallel orientation. The pathways are represented as an oriented graph where the nodes are the knot or link types, and two nodes are connected by an edge if one can go from one to the other via a reconnection event. The substrate and product of each reconnection are indicated by the orientation of the edges. The diagrams above each edge illustrate an example of the corresponding reconnection event by showing the band where the band surgery will be performed. The weights on the edges correspond to transition probabilities obtained numerically. Details of the simulations are in the Numerical Methods section below, and in the Supplementary Methods and Supplementary Data online.

The 9 pathways found in Theorem 2 involve 16 possible transitions taking a knot to a link or vice versa; 6 of the transitions have fully characterized mechanisms. The proof of the theorem and the characterization of the mechanisms are presented in the Supplementary Methods online. Fig. 4 summarizes the results as an oriented graph where each node is a knot/link type and each edge represents the transition between two topologies by one reconnection step. All minimal pathways taking the parallel 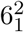 to the unlink 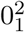, and satisfying Assumption 1 are shown. In the next section we undertake a thorough computational study with the objective of discriminating between minimal pathways while minimizing the number of assumptions. In particular, we use the numerical work to assign frequencies to each transition in the pathway graph (represented in Fig. 4 as weights on the edges).

We here give a draft of the proof of Theorem 2. More details, including Lemmas S1-S8, Propositions S9-S17, and Figs. S1 and S2 exhibiting the steps of the proof and relevant band surgeries for each of the transitions in Fig. 4, are included in the Supplementary Methods online. In order to characterize the minimal pathways starting from the parallel 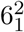 link, we first investigate the effect of band surgeries on certain topological invariants such as the signature, the Jones polynomial, the Q polynomial and the Arf invariant of the knots and links involved in those pathways. By Lemma S6, the sequence of the signatures of knots and links is –5, –4, –3, –2, –1,0,0. Lemma S7 shows that split links can not appear in a shortest pathways. Lemma S8 identifies the candidate topologies for the minimal pathways from 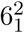

#### Outline of the proof

(First step) From Proposition S9, the product knot obtained from 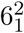 is either 5_1_ or 3_1_#3_1_.

(Second step) From Proposition S10, the product link obtained from 5_1_ is either 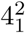 or 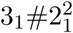. From Proposition S11, the product link obtained from 3_1_#3_1_ is either 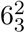 or 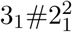.

(Third step) From Proposition S12, the product knot obtained from 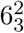 is 5_2_. From Proposition S13, the product knot obtained from 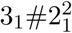 is either 5_2_ or 3_1_. From Proposition S14, the product knot obtained from 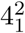 is 3_1_.

(Fourth step) From Proposition S15, the product link obtained from 5_2_ is either 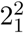 or 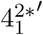. From Proposition S16, the product link obtained from 3_1_ is 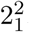.

(Fifth step) From Proposition S17, the product knot obtained from 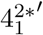 is 0_1_. The product obtained from 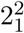 is 0_1_. In the last step, the recombination event changes 0_1_ into 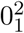. These steps cover all transitions satisfying the Assumption 1.

### 2.2 Topological mechanisms of reconnection

The topological mechanisms of events between the following (substrate, product) pairs have been fully characterized:^7^ 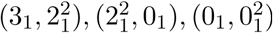. The topological mechanisms between pairs 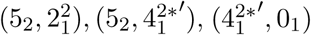 are characterized in the proposition below. For all transitions along the 9 minimal pathways, Fig. 4 illustrates one possible band surgery relating the knot to the link. The proof of Proposition 3 is given in the Supplementary Methods online, Characterization of Mechanisms section (Supplementary Fig. S3, Proposition S18, Theorem S19, Lemma S20).

#### Proposition 3.

A.^25^ Suppose 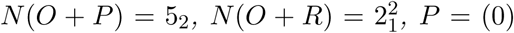 and *R* =(*w*, 0).Then 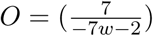

B.^25^ Suppose 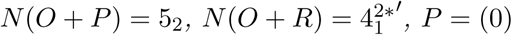 and *R* =(*w*, 0).Then 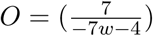.

C.^26^ Suppose 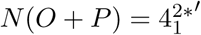, *N*(*O* + *R*) = 01,*P*= (0)and *R* =(*w*, 0).Then 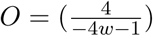. Because XerC and XerD are tyrosine recombinases and act through a Holliday Junction Intermediate, the tangle pairs (*P*, *R*) that are relevant to unlinking of DNA replication links by Xer recombination are (*P*, *R*) = ((0)_*p*_, (–1)), (*P*,*R*) = ((0)_*a*_, (0,0)) (*P*, *R*) = ((0)_*p*_, (1)) as illustrated in Fig. 1C. The above proposition allows to determine all the topological mechanisms for each of the three combinations of substrate and product in the statement. We illustrate the solutions in Proposition S18 and in Supplementary Fig. S3 in the Supplementary Methods online. Just as in Shimokawa *et al.*,^7^ here each system of tangle equations yields three solutions, and the three solutions can be interpreted as representing a unique 3-dimensional topological mechanism.

### 2.3 Which unlinking pathways are most probable?

In the previous section, we proved analytically that under Assumption 1 there are 9 minimal pathways of unlinking the parallel 6-cat, 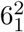. The mathematical analysis that includes enumeration of pathways and characterization of topological mechanisms becomes difficult for substrates with high crossing numbers. Furthermore, if the assumption of reduction in complexity ‐which is equivalent to imposing a topological filter in the physical system‐ is lifted, then the number of possible pathways increases rapidly and the detailed mathematical analysis quickly becomes intractable. We here remove Assumption 1 and set out on a numerical exploration of reconnection pathways starting from a broader set of substrate topologies. We develop software which finds reconnection sites along polygonal chains in the simple cubic lattice and simulates the reconnection event. Fig. 5C illustrates the basic reconnection move on a simplified polygon. Fig. 5A shows a lattice trefoil with one single reconnection site, before and after local reconnection. We simulate reconnection to explore different topological transitions, to quantify transition probabilities and to discriminate between unlinking pathways that are mathematically indistinguishable when only substrate, product and length are specified.

**Figure 5:**
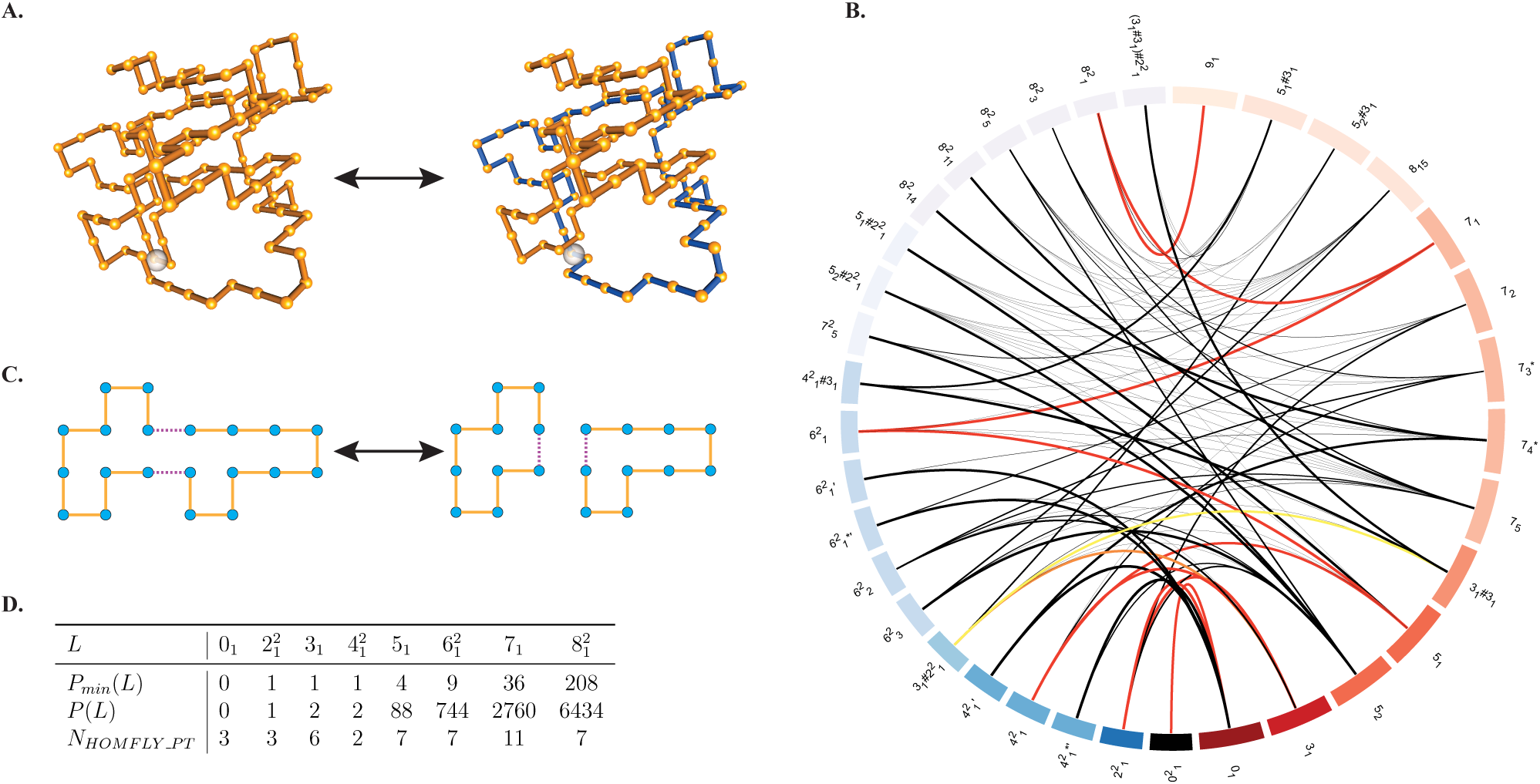
(A) The substrate (left) is a lattice trefoil with 120 segments and two directly repeated reconnection sites indicated by a white sphere. The product (right) is a 2-component link obtained after one reconnection event. All substrate knots have directly repeated sites that are 60 segments apart, with a tolerance of ±6 segments, and all links have two components 60 ± 6 long so that the sum of the lengths is exactly 120. Reconnection on links is only performed between sites in different components. (B) Circos figure: all reconnection transitions in a minimal pathway from the 9_1_ that satisfy Assumption1. 2-component links (resp. knots) are arranged by increasing crossing number from bottom to top in the left (resp. right) hemisphere, and are color-coded blue (resp. red). Color intensity increases with decreasing crossing number. An arc between *K* and *L* indicates at least one observed reconnection event between *K* and *L*. The thickness of the arcs corresponds to the directed transition probability between two topologies. Transitions with an observed probability < .2 are thickened to be more visible. Transitions are colored according to the probability of the most probable minimal pathway they are a member of. The first, second, and third most probable unlinking pathways from 9_1_ are colored red, orange, and yellow, respectively. If no arc appears between a pair {*K*, *L*}, this means that no reconnection between them was observed. Observed transitions for all substrate topologies, including those in non-minimal pathways, are included in Supplementary Data and in Fig. S6 in the Supplementary Methods online. (C) Local reconnection move between two directly repeated sites. In the juxtaposition the reconnection sites, indicated with hashed lines, are at distance 1 and in antiparallel alignment. (D) *L* are *T*(2, *n*) torus knots and links (Fig.1). *P*_*min*_(*L*) is the number of minimal unlinking pathways observed for *L* under Assumption1. *P*(*L*) indicates the total number of minimal pathways observed for *L* without Assumption! It is known that there are infinitely many minimal unlinking pathways for any *T*(2, 2*n*) link with parallel sites.^17^ Nhomfly-pt is the number of distinct HOMFLY-PT polynomials observed after one reconnection.

We provide numerical evidence that, of all minimal pathways starting with the RH parallel 6-cat, the one in Fig. 1A is the most likely. The weights in Fig. 4 correspond to the transition probabilities obtained in the numerical simulations. More generally, our numerical data suggest that this trend holds for any substrate that is a RH 2*m*-cat with parallel sites, or a RH (2*m* – 1)-torus knot with two sites in direct repeats. It is important to emphasize that the simulations do not use Assumption 1. Fig. 5B is a circos figure that shows all observed reconnection transitions that maintain or decrease minimal crossing number and that belong to an observed minimal pathway from the9_1_knot. The thickness of the arcs corresponds to the directed transition probability between two topologies. Transitions in the most probable minimal pathway from 9_1_ are colored red. The predominance of these most probable unlinking pathways is consistent with the experimental observations for XerCD-FtsK-*dif* site-specific recombination on DNA replication links,^10^ and for reconnection in fluid vortices,^12^ and is also consistent with the predictions in the literature.^7,11^

The minimum distance between the link type *L*_*i*_ and the knot type *K*_*j*_ in terms of band surgeries is called *nullification distance*.^27,28^ In the numerical experiment we started by choosing knots and 2-component links that are at nullification distance 1-3 from one of the 11 knots or links along one of the 9 minimal pathways of Theorem 2 and Fig. 4, or are obtained from these topologies by taking mirror images or reversing the orientation of one of the components. For completeness, we expanded the initial set to include 491 substrate topologies representing almost all knots and links with 9 or fewer crossings. Reasons for omitting a handful of 9-crossing split links from the substrate set are described in detail below. We use the BFACF algorithm to generate large independent ensembles of conformations for each substrate topology. BFACF is a dynamic Monte Carlo method which samples uniformly the set of all lattice polygons of fixed topology for a given mean length.^29^ The BFACF moves used to perturb each chain are illustrated in Fig. S4 in the Supplementary Methods online. Split links such as the unlink 01 or 0_1_U3_1_ (see Fig. 3), even though they appear as reconnection products, are not used as substrates due to the difficulty of keeping the components together without altering the Monte Carlo procedure. In order to improve the efficiency of sampling statistically independent conformations we implemented BFACF as a Composite Markov Chain (CMC). Details of the simulations, including a description of the algorithms and different parameters, are included in the numerical methods section and in the Supplementary Methods. Fig. S6 in the Supplementary Methods online illustrates all the transitions observed between 881 topologies in the numerical experiment, including those that do not appear in minimal pathways from 9_1_. The resulting transition probabilities are available in matrix form in the data spreadsheet provided as Supplementary Information (Supplementary Data).

Fig. 5D contains exact counts for the number of minimal unlinking pathways for torus knots and links with up to 6 crossings, and the corresponding numerical estimates for 7 and 8 crossings. Under Assumption 1 there are 9 minimal pathways of unlinking the 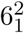 link. In the numerical study, we find 36 minimal unlinking pathways for the 7_1_ knot and 208 minimal unlinking pathways for the 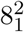 link, under Assumption 1 (*P*_*min*_(*L*)). Once the Assumption is removed, we observe *P*(7_1_) = 2760 minimal pathways for the knot 7_1_ and 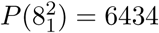 minimal pathways for the link 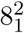 (in this case the crossing number can increase at any given step). However it has been shown analytically that there are infinitely many possible minimal pathways between any 2*n* torus link with parallel sites and the unlink.^17^ The numerical data can provide biologically-relevant information by establishing a ranking of the most likely pathways. The third row in Fig. 5D indicates the number of distinct product topologies (as detected by the HOMFLY-PT polynomial) observed for torus knots and links of the type *T*(2,*n*) with 8 or fewer crossings after a single reconnection step.

## 3 Discussion

In Theorem 2 we prove that there are exactly 9 shortest unlinking pathways for the 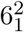, assuming that at every step the complexity of the substrate goes down or remains the same. The 9 pathways are illustrated in Fig. 4. We solve the topological mechanisms involved for 6 of the 16 steps along these pathways. We develop a new Monte Carlo based numerical method which allows us to model local reconnection on chains of fixed length and topology. We run the numerical simulation on each topology found to be within 3 nullification steps from any topology in Fig. 4. Notice that in these experiments there is nothing preventing the complexity of a substrate from going up at any given step. We can determine the set of all minimal pathways from any of the substrate topologies, and single out the most probable pathway. In Fig. 5 we provide numerical estimates for the number of minimal pathways for torus knots and links with 7 and 8 crossings. In our numerical data the most probable minimal pathway from a torus link (or knot) to the unlink is the one where every intermediate is also in the torus family as in Fig. 1A. The data from the numerical experiments can be found in the Supplementary Data.

Mathematically, extending Theorem 2 to determine all minimal pathways for *T*(2, *N*) torus knots and links is difficult. In general, if the substrate is a torus knot or link *T*(2,*N*) one can find multiple pathways that preserve the minimal crossing number at many steps. The complexity of the problem grows with the minimal crossing number of the substrate. For example, using numerical simulation we estimate the number of minimal pathways from the 7_1_ (resp. 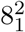) to the unlink to be at least 36 (resp. 208) under Assumption 1. These are not tight bounds due to the limitations with using links of the form 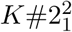 as substrates in the numerical experiments. It is known that when the assumption is removed, there are infinitely many shortest pathways between the *T*(2, 2*N*)_*p*_ torus link and the unlink.^17^ In our numerical work, once Assumption 1 is removed we count at least 744, 2760 and 6434 shortest unlinking pathways for 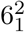, 7_1_ and 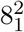, respectively.

The problem of computing the nullification distance between a knot and a link is of interest to the mathematical community.^17,23,24,27,28^ In cases where the analytical tools fail to provide an exact nullification distance, one can estimate the distance between two topologies using the numerical method and possibly remove ambiguities by exhibiting the relevant band surgeries.

The numerical simulations in this study posed a number of challenges. For example, in order to generate an ensemble of essentially independent unknots 0_1_ of length 120 we had to go through at least twice as many iterations of the BFACF algorithm than for any other substrate topology. Further, these unknots contained synapses meeting the reconnection criteria approximately once every 7.5 × 10^9^ iterations. In order to improve the efficiency of such runs, we implemented the BFACF algorithm as a Composite Markov Chain process.^30–33^ Similar challenges extend to any topology consisting of a connected sum of a knot and a Hopf link 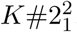, or the disjoint union of a knot and an unknot *K* ⋃ 0_1_ (see examples in Fig. 3). In the first case, the unknotted component tends to shrink, making it difficult to satisfy the equal-length criteria for recombination. In the second case, even though these topologies appear as reconnection products, they cannot be used as substrates due to the difficulty of keeping the components together (*w*ithout biasing the simulations for those specific substrates). Now consider an example where a bacterial chromosome dimer forms a 31 knot with two equidistant directly repeated *dif* sites. In our simulations we see that 0.025% of trefoils transition to 0_1_ ⋃ 3_1_, the disjoint union of an unknot and a trefoil, and 95.2% of trefoils transition to 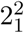. In the first case the knotted dimer is effectively unlinked in one step, but one of the components will remain knotted, which can pose problems during chromosome segregation. In the second case unlinking of the trefoil can be achieved in 3 steps, with a combined probability of 0.925; the final product is 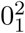, a union of two circles which can then segregate at cell division.

In the case of unlinking of DNA replication links, each component of the link corresponds to a newly replicated chromosome from *E.coli* with one *dif* site in each component. This example motivated our choice to let two reconnection sites within a single circle be equidistant, and the two components of a linked product or substrate have the same length. In different contexts, such as that of site-specific recombination between non-equidistant sites, more general homologous recombination, and possibly other reconnections in physics, the distance between sites will be an important parameter, requiring further exploration of the length and topology dependence of the transition probabilities obtained by the numerical method.

Furthermore, in nature, DNA molecules are often found tightly packaged in crowded environments. A study of reconnection on confined chains would shed light on whether confinement plays a role in driving topological simplification by any process involving local reconnection. Existing studies of the confinement of polygonal chains inside and outside the lattice suggest methods for generating ensembles of conformations.^34,35^

## 4 Materials and Methods

### 4.1 Mathematical Methods

The tangle method is briefly summarized in Fig. 1. The naming convention used for knots and links is reviewed in the Introduction. More detailed mathematical methods and results used in the proof of Theorem 2 are provided in Fig. 4 and in the Supplementary Methods online. A site-specific recombination event is modeled as a local reconnection and is represented mathematically as a system of tangle equations as described in Fig. 1B. The circular chain represents the starting knot or link, and *P* is a 2-string tangle that encloses the reconnection sites. Reconnection changes *P* into *R*. We assume that each reconnection is modeled as a coherent band surgery, i.e. *P* = (0) and *R* = (*w*; 0) for some integer *w* (Fig. 1C).

### 4.2 Numerical Methods: modeling reconnection

#### Computer simulations of local reconnection

We use an integrated set of computational tools to generate and filter ensembles of conformations, perform reconnection, identify product topologies, generate transition probabilities and facilitate statistical analysis of the results. Given an ensemble of lattice conformations with fixed length and constant topology, our algorithm searches for possible synapses along each conformation, selects one uniformly at random, and performs reconnection as illustrated in Fig. 5A. Our original motivation came from XerC/D site-specific recombination at *dif* sites in newly replicated chromosomes with one site in each component or in chromosome dimers with two equidistant directly-repeated sites. In this case reconnection events are constrained by the position and orientation of the *dif* sites. We therefore impose a set of constraints on where to perform reconnection. These can be seen as topological filters that can be adjusted to best fit the scenario to be modeled. Here, a reconnection synapse is defined as a pair of coplanar edges of distance one apart with antiparallel orientation; each of the two oriented edges is a *reconnection site*. Reconnection exchanges each edge of the synapse for one perpendicular to it as shown in Fig. 5C. The set of possible edge pairs on which to form a synapse is further constrained by step distance along the conformation. Here we adjust this parameter to constrain the location of the synapse so that the arc lengths on each side are equal within a ±6 range, while enforcing the total length of the knotted polygon, or the sum of the lengths of the components of interlinked polygons, to be fixed. For knots this models two equidistant sites in the synapse. For two component links, it models two components of equal length with a single site in each of the two components. We exclusively sampled conformations of total length 120 which contain at least one reconnection synapse.

#### Generation of reconnection substrates

Self-avoidance is an important property when modeling biopolymers such as circular DNA. Here, conformations in the simple cubic lattice, ℤ^3^, are self-avoiding polygons whose vertices have integer coordinates and whose edges are parallel to one of the three coordinate axes. The BFACF algorithm is a dynamic Monte Carlo method which samples from the space of lattice conformations of a fixed topology.^29^ The states of the resulting Markov Chain are conformations obtained by first randomly selecting an edge, then attempting one of the three moves shown in Fig. S4 in the Supplementary Methods online ((-2)-move, (+2)-move or (0)-move). None of these moves can ever change the link type of the conformation.^29,36^

Generating large ensembles of conformations for each topology with at least one valid synapse posed significant technical challenges. The 0_1_ knot and links of the type 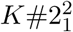 where *K* is a knot with high crossing number were particularly problematic. This is because the component with trivial topology tends to have a short average length, making sampled conformations that form a reconnection synapse very rare. For example, the 0_1_ forms such a synapse in fewer than 1 in 1.3 × 10^6^ sampled conformations. To address these challenges and gain the computational performance needed for this study, we here extend the efficient, constant time (in knot length) implementation of the BFACF algorithm used in previous work^34,35,37,38^ by employing it as a Composite Markov Chain (CMC) Monte Carlo process.^30–33,39^ CMC BFACF iterates simultaneously on multiple Markov chains with different fugacity parameters, swapping conformations between chains when certain weighted random criteria are met; more details of the implementation are included in the Supplementary Methods online. CMC Monte Carlo improves efficiency by exchanging conformational states between chains, thus improving the speed at which the conformations are randomized. We sample conformations at a frequent fixed rate and correct for dependent samples using block mean analysis,^40^ therefore standardizing the sampling methodology across all of the topologies in the study and avoiding reliance on direct estimations of integrated autocorrelation time. With this methodology, we generated in the range of 10^7^ conformations for every substrate topology. Of the topologies for which a reconnection event was observed, the number of conformations containing at least one reconnection synapse ranged from approximately 1.5 × 10^6^ for the 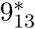 knot, to as little as 86 for the 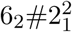 link. Two component topologies in which the two components are of different topology are difficult to sample efficiently because of the rarity of conformations that meet our stringent arclength criteria. Split links, *i.e.* those topologies in which the two components are not interlinked, are even more problematic because both components tend to travel away from each other, thus dramatically reducing the probability of sampling conformations that contain a valid synapse. We identified those topologies as products of reconnection, but did not include them in the set of substrate topologies described in the next paragraph.

Recall that 9 minimal unlinking pathways from the 6-cat were obtained analytically in Theorem 2 under the assumption that each reconnection step either preserves or reduces the complexity of the l· substrate. Our simulations eliminate that assumption, enabling wider exploration of possible topo; logical reconnection pathways. We start with 491 substrate topologies, including those along the 9 unique pathways from Fig. 4 (excluding the unlink 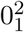). With CMC BFACF we generate ensembles of conformations with fixed topology to be used as reconnection substrates. The number of substrate conformations generated ranges from 1.2 × 10^7^ for the 72 link, to more than 6.9 × 10^8^ for the 0_1_. We perform one reconnection per conformation and identify the resulting topology. Including all substrate topologies and the identifiable products after reconnection, there are 881 topologies being analyzed in the study (490 knots and 391 two component links).

#### Knot identification

Our simulations require a rigorous, unambiguous way of identifying the knot or link conformation l· types in ℤ^3^. With the exception of chiral knots 8_17_ and 9_42_ which have the same HOMFLY-PT as their mirror images, and 9_12_ which has the same HOMFLY-PT as 4_1_#5_2_, all prime knots with nine or fewer crossings can be unambiguously identified using the HOMFLY-PT polynomial.^41,42^ Our knot
-identification software is based on the other published algorithms.^43,44^ In order to identify product topologies, we first perform 20,000 BFACF iterations with randomly chosen (0) and (-2) moves. At each step, the conformation either remains the same length or becomes shorter, in many cases approaching the minimal length for that topology.^38^ The final conformation goes through an energy minimization algorithm,^22^ we compute an extended Gauss code and identify the topology using the HOMFLY-PT polynomial. Information on those oriented knots or links with 10 or fewer crossings that HOMFLY-PT fails to identify uniquely is included in the Supplementary Methods online.

Recombination between two directly repeated sites along a single circular chain yields a 2-component link. The number of product topologies increases dramatically with the complexity of the substrate. Fig. 3 shows a selection of some of the expected products, including composite links that are not-normally shown in knot tables. Composites are of two types: connected sums of prime knots or links; and disjoint unions. In this study, we perform recombination on two types of substrates: (i) knots with two (approximately) equidistant directly repeated sites; and (ii) links with 2 components of identical total length and with one site in each component. More specifically, each substrate knot is a self-avoiding lattice polygon of length 120 and recombination occurs on two directly repeated sites that are between 54 and 66 units apart (Fig. 5A). Each linked substrate consists of two self-avoiding polygons between 54 and 66 units long, such that the sum of their lengths is exactly 120. Recombination is restricted to synapses where two sites, one in each component, are found at unit distance apart and in anti-parallel alignment as illustrated in Fig. 5(A and C). A small representative subset of the knot and link types used in the simulations is shown in Fig. 3, and the naming convention is described in the nomenclature section, in the Supplementary Methods and in Supplementary Fig. S5 online.

## Acknowledgments

This research was supported by the following: Japan Society for the Promotion of Science KAKENHI grant numbers 25400080, 26310206, 16H03928, 16K13751, 17H06463(to K.S.), 26800081 (to K.I.); National Science Foundation DMS1716987 (MF, MV) and CAREER Grant DMS1057284 (MV, RS, MF, RB) and NIH-R01GM109457 (MV); Welcome Trust SIA 099204/Z/12Z and 200782/Z/16/Z (DJS). The authors are grateful to R. Scharein for providing assistance with Knotplot and for his work on the first version of the reconnection software; C. Soteros, M. Szafron and M. Schmirler for contributing their statistical expertise; J. Arsuaga, D.W. Sumners and S. Witte for helpful discussions; and Barbara Ustanko, ELS, for editorial assistance with this manuscript.

## 6 Author contributions

MV conceived the overall research project. MV, KS and DS conceived the detailed research plan. MV and KS directed the mathematical component of the paper. MV and RB directed the computational component of the paper. MY and KI performed the details of the mathematical research. RS, MF and RB performed the details of the computational component. MV, KS wrote the main manuscript text; MV, RB and RS wrote the numerical methods; MV, KS, KI wrote the mathematical methods and proofs. RS, KI, KS, MF, MV and MY prepared figures for publication. All authors reviewed the manuscript.

## References

1 Navashin, M. S. Unbalanced somatic chromosomal variation in Crepis. Univ. Calif. Pub. Agr. Sci. 6, 95–106 (1930).

2 McClintock, B. A correlation of ring-shaped chromosomes with variation in Zea Mays. Proc. Natl. Acad. Sci. USA 18, 677–681. (1932).

3 Wang, J. C. & Schwartz, H. Noncomplementarity in base sequences between the cohesive ends of coliphages 186 and lambda and the formation of interlocked rings between the two DNA’s. Biopolymers 5, 953–966 (1967).

4 Sundin, O. & Varshavsky, A. Terminal stages of SV40 DNA replication proceed via multiply inter-twined catenated dimers. Cell 21, 103–114 (1980).

5 Wasserman, S. & Cozzarelli, N. Biochemical topology: applications to DNA recombination and replication. Science 232, 951–960 (1986).

6 Adams, D. E., Shekhtman, E. M., Zechiedrich, E. L., Schmid, M. B. & Cozzarelli, N. R. The role of topoisomerase IV in partitioning bacterial replicons and the structure of catenated intermediates in DNA replication. Cell 71, 277–288 (1992).

7 Shimokawa, K., Ishihara, K., Grainge, I., Sherratt, D. J. & Vazquez, M. FtsK-dependent XerCD-dif recombination unlinks replication catenanes in a stepwise manner. Proc. Natl. Acad. Sci. USA 110, 20906–20911 (2013).

8 Sogo, J., Greenstein, M. & Skalka, A. The circle mode of replication of bacteriophage lambda: the role of covalently closed templates and the formation of mixed catenated dimers. J. Mol. Biol. 103, 537–562 (1976).

9 Zechiedrich, E. L., Khodursky, A. B. & Cozzarelli, N. R. Topoisomerase IV, not gyrase, decatenates products of site-specific recombination in Escherichia coli. Genes Dev. 11, 2580–2592 (1997).

10 Grainge, I. et al. Unlinking chromosomes catenated in vivo by site-specific recombination. EMBO J. 26, 4228–4238 (2007).

11 Kleckner, D., Kauffman, L. H. & Irvine, W. T. M. How superfluid vortex knots untie. Nat. Phys. 12, 650–655 (2016).

12 Kleckner, D. & Irvine, W. T. M. Creation and dynamics of knotted vortices. Nat. Phys. 9, 253–258 (2013).

13 Laing, C. E., Ricca, R. L. & Sumners, D. W. L. Conservation of writhe helicity under anti-parallel reconnection. Scientific Reports 5, 9224 10.1038/srep09224 (2015).

14 Ishihara, K. & Shimokawa, K. Band surgeries between knots and links with small crossing numbers. Prog. Theor. Phys. Supplement 191, 245–255 (2011).

15 Ishihara, K., Shimokawa, K. & Vazquez, M. Site-specific recombination modeled as a band surgery: applications to Xer recombination In: Jonoska, N., Saito, M. (eds) Discrete and topological models in molecular biology, 387–401. Nat. Comput. (Springer, Heidelberg, Berlin, Heidelberg, 2014).

16 Yoshida, M. Applications of band surgery and signed crossing changes of knots and links to molecular biology. Master’s thesis, Department of Mathematics, Saitama University (2013).

17 Buck, D. & Ishihara, K. Coherent band pathways between knots and links. J. Knot Theory Ramifications 24, 1550006–27 (2015).

18 Buck, D., Ishihara, K., Rathbun, M. & Shimokawa, K. Band surgeries and crossing changes between fibered links. J. London Math. Soc. 94, 557–582 (2016).

19 Ip, S. C. Y., Bregu, M., Barre, F.-X. & Sherratt, D. J. Decatenation of DNA circles by FtsK-dependent Xer site-specific recombination. EMBO J. 22, 6399–6407 (2003).

20 Rolfsen, D. Knots and links (AMS Chelsea, Providence, RI, 2003).

21 Brasher, R., Scharein, R. G. & Vazquez, M. New biologically motivated knot table. Biochem Soc. Trans. 41, 606–611 (2013).

22 Scharein, R. G. Interactive topological drawing. Ph.D. thesis, Department of Computer Science, The University of British Columbia (1998).

23 Kanenobu, T. Band surgery on knots and links. J. Knot Theory Ramifications 19, 1535–1547 (2010).

24 Kanenobu, T. Band surgery on knots and links, II. J. Knot Theory Ramifications 21, 1250086–108 (2012).

25 Darcy, I. K., Ishihara, K., Medikonduri, R. K. & Shimokawa, K. Rational tangle surgery and Xer recombination on catenanes. Algebr. Geom. Topol. 12, 1183–1210 (2012).

26 Vazquez, M., Colloms, S. & Sumners, D. Tangle analysis of Xer recombination reveals only three solutions, all consistent with a single three-dimensional topological pathway. J. Mol. Biol. 346, 493–504 (2005).

27 Diao, Y., Ernst, C. & Montemayor, A. Nullification of knots and links. J. Knot Theory Ramifications 21, 1250046–70 (2012).

28 Ernst, C. & Montemayor, A. Nullification of torus knots and links. J. Knot Theory Ramifications 23, 1450058–77 (2014).

29 Madras, N. & Slade, G. The Self-Avoiding Walk (Modern Birkhäuser Classics, Cambridge, MA, 1996).

30 Geyer, C. J. Practical Markov chain Monte Carlo. Statistical Science 7, 473–483 (1992).

31 Orlandini, E. Monte Carlo Study of Polymer Systems by Multiple Markov Chain Method, in Numerical Methods for Polymeric Systems, 33–57 (Springer New York, New York, NY, 1998).

32 Szafron, M. Monte Carlo Simulations of Strand Passage in Unknotted Self-Avoiding Polygons. Master’s thesis, Department of Mathematics and Statistics, University of Saskatchewan (2000).

33 Szafron, M. Knotting statistics after a local strand passage in unknotted self-avoiding polygons in Z3. Ph.D. thesis, Department of Mathematics and Statistics, University of Saskatchewan (2009).

34 Ishihara, K. et al. Bounds for the minimum step number of knots confined to slabs in the simple cubic lattice. J. Phys. A: Math. Theor. 45, 065003–27 (2012).

35 Arsuaga, J. et al. Current theoretical models fail to predict the topological complexity of the human genome. Front. Mol. Biosci. 2, 48 (2015).

36 Janse van Rensburg, E. J., Orlandini, E., Sumners, M. C. D w and. Tesi & Whittington, S. G. The writhe of knots in the cubic lattice. J. Knot Theory Ramifications 6, 31–44 (1997).

37 Hua, X., Nguyen, D., Raghavan, B., Arsuaga, J. & Vazquez, M. Random state transitions of knots: a first step towards modeling unknotting by type II topoisomerases. Topol. Appl. 154, 1381–1397 (2007).

38 Scharein, R. et al. Bounds for the minimum step number of knots in the simple cubic lattice. J. Phys. A: Math. Theor. 42, 475006 (2009).

39 Orlandini, E., Janse van Rensburg, E. J., Tesi, M. C. & Whittington, S. G. Entropic Exponents of Knotted Lattice Polygons, in Topology and Geometry in Polymer Science, vol. 103 (Springer, Berlin, 1998).

40 Fishman, G. Discrete-event simulation: modeling, programming, and analysis (Springer-Verlag, London, 2001).

41 Freyd, P. et al. A new polynomial invariant of knots and links. Bull. Amer. Math. Soc. 12, 239–246 (1985).

42 Przytycki, J. H. & Traczyk, P. Conway algebras and skein equivalence of links. Proc. Amer. Math. Soc. 100, 744–748 (1987).

43 Gouesbet, G., Meunier-Guttin-Cluzel, S. & Letellier, C. Computer evaluation of homfly polynomials by using gauss codes, with a skein-template algorithm. Appl. Math. Comput. 105, 271–289 (1999).

44 Jenkins, R. J. Knot Theory, Simple Weaves, and an Algorithm for Computing the HOMFLY Polynomial. Master’s thesis, Carnegie Mellon University (1989).

